# Advancing Liver Gene Therapy: Enhanced Transduction with GalNAc-Bioconjugated rAAV Capsids

**DOI:** 10.1101/2024.07.25.605063

**Authors:** Pierre-Alban Lalys, Audrey Bourdon, Mohammed Bouzelha, Dimitri Alvarez-Dorta, Karine Pavageau, Tiphaine Girard, Roxane Peumery, Zakaria Bouchouireb, Maia Marchand, Anthony Mellet, Mireille Ledevin, Sébastien Depienne, Mickaël Guilbaud, Mikaël Croyal, Sébastien G. Gouin, Benoit Roubinet, Ludovic Landemarre, Oumeya Adjali, Eduard Ayuso, Thibaut Larcher, Caroline Le Guiner, Mathieu Mével, David Deniaud

**Affiliations:** Nantes Université, CNRS, CEISAM UMR 6230, F-44000 Nantes, France; Nantes Université, CHU de Nantes, INSERM UMR 1089, Translational Gene Therapy Laboratory, F-44200 Nantes, France; Capacités, 16 rue des marchandises, 44200 Nantes, France; INRAE, Oniris, PanTher, APEX, F-44307 Nantes, France; Nantes Université, CNRS, INSERM, Institut du thorax, F-44000 Nantes, France; Nantes Université, CHU Nantes, Inserm, CNRS, SFR Santé, Inserm UMS 016, CNRS UMS 3556, F-44000 Nantes, France; Glycodiag, 2 rue du cristal, 45100 Orléans, France

**Keywords:** adeno-associated virus, chemistry, bioconjugation, gene therapy, liver

## Abstract

This study investigates novel approaches to improve targeted gene delivery to the liver, a crucial organ for metabolic processes that faces vulnerabilities from various pathologies. Adeno-associated virus (AAV)-based gene therapy has emerged as a promising approach for liver targeting, with numerous investigational avenues. However, administration of high doses of AAV vectors present safety concerns, often requiring the use of corticosteroids and immunosuppression to mitigate immune adverse events. To address this, substantial efforts are underway to engineer optimized capsids to enhance the efficiency and specificity of recombinant AAV (rAAV) targeting hepatocytes, aiming to reduce required dosages. In this study, we employed bioconjugation chemistry to target the Asialoglycoprotein receptor (ASGPR), a C-type lectin abundantly expressed at the surface of hepatocyte membranes. We demonstrated that covalently attaching carbohydrates derived from GalNAc (a known ASGPR ligand) to lysine amino-acids on the rAAV2 capsid significantly enhanced *in vivo* liver transduction efficiency in mice. These optimized vectors present a promising avenue for the treatment of a spectrum of liver diseases, providing an alternative solution within the framework of liver gene therapy.

## Introduction

The liver, with its myriad functions in nutrient and xenobiotic metabolism, plays a pivotal role in maintaining overall health. The orchestration of these intricate tasks relies on various proteins, and mutations in their encoding genes can lead to diseases or predisposition to them. Conditions associated with these mutations encompass defects in carbohydrate and lipid metabolism, bleeding disorders, increased susceptibility to drugs and toxins, and cognitive deficits.[1,2]

Given the essential role of the liver, the development of strategies to improve liver-targeted delivery of therapeutics is of significance importance. The ability to introduce functional gene expression cassettes into hepatocytes, efficiently compensating for primary liver defects, represents a potent therapeutic tool.[3] Moreover, the sophisticated secretory apparatus of the liver provides an opportunity to express and release proteins not typically synthesized in this organ. This capability extends to the treatment of diseases or deficiencies such as hemophilia,[4] Wilson’s disease,[5] alpha-1 antitrypsin deficiency,[6] or hemochromatosis.[7]

Among current treatments, viral vector-based gene therapy can be mentioned. Indeed, clinical trials utilizing recombinant adeno-associated virus (rAAV)-mediated gene therapy for liver-focused metabolic and genetic diseases have shown promising results. Notably, the effectiveness of rAAV gene therapy in treating individuals with hemophilia has gained increased attention, as evidenced by the recent approvals of valoctocogene roxaparvovec (Roctavian™) for Hemophilia A [8], etranacogene dezaparvovec (Hemgenix®) for Hemophilia B [9] by both the EMA (European Medicines Agency) and the FDA (Food and Drug Administration), and fidanacogene elaparvovec-dzkt (Beqvez®) for Hemophilia B [10] by the FDA.

Despite these successes, challenges related to vector toxicity and immunogenicity persist, particularly in instances requiring high vector doses.[11] Consequently, there is a clear to enhance the safety and efficacy of recombinant AAV vectors, prompting the exploration of advanced targeting strategies for gene therapy.[12] A precise and efficient delivery to specific tissues or cell types would minimizes adverse off-target effects and provides an opportunity to reduce the injected dose required for patients.

Hepatocytes, expressing a high density of asialoglycoprotein receptor (ASGPR), present a strategic target for achieving selective and efficient gene delivery to the liver.[13] Indeed, this receptor is a carbohydrate-binding protein that recognizes and binds *N*-acetylgalactosamine (GalNAc) or galactose residues.[14,15] By engineering rAAV vectors to recognize and bind to ASGPR, the specificity of gene expression in hepatocytes could be enhanced.[16,17] This targeted approach holds great potential to treat genetic or metabolic disorders in the liver, with improved therapeutic precision and safety.

In our study, we designed a range of bifunctional ASGPR-binding ligands having a reactive bioconjugation function to chemically functionalize the rAAV2 capsid. We demonstrated that chemically modifying the amino group of the lysine residues of the rAAV2 capsid with GalNAc-derived carbohydrates resulted in a substantial enhancement in hepatocyte transduction *in vitro* and *in vivo* as compared to unmodified rAAV2. This direct bioconjugation strategy holds significant potential for fine-tuning rAAV capsid immunoreactivity and tropism across various therapeutic applications.

## Materials and methods

The synthesis and the characterizations of the compounds 1-4 are detailed in the supporting information part (**Figure S-1 – S-6**).

### -rAAV2-CAG-GFP production and purification

AAV2 vectors were produced from two plasmids: (i) pHelper, PDP2-KANA encoding AAV Rep2-Cap2 and adenovirus helper genes (E2A, VA RNA, and E4); and (ii) the pVector ss-CAG-eGFP containing the ITRs. All vectors were produced by transient transfection of HEK293 cells using the calcium phosphate-HeBS method. AAV2 transfected cells were harvested 48 h after transfection and treated with Triton-1% and benzonase (25 U/mL) for 1 h at 37°C. The resulting bulk was subjected to freeze-thaw cycles to release vector particles. The cellular debris were removed by centrifugation at 2500 rpm for 15 min. Cell lysates were precipitated with PEG overnight and clarified by centrifugation at 4000 rpm for 1 h.

The precipitates were then incubated with benzonase for 30 min at 37 °C and collected after centrifugation at 10,000 g for 10 min at 4°C. Vectors were purified by double cesium chloride (CsCl) gradient ultracentrifugation. The viral suspension was then subjected to four successive rounds of dialysis with mild stirring in a Slide-a-Lyzer cassette (Pierce) against dPBS (containing Ca++ and Mg++).

### Bioconjugation and purification

rAAV2-CAG-GFP (1E12 vg, 2.49 nmol) were added to a solution of TBS buffer (pH 9.3) containing compounds **1**, **2**, **3** or **4** (3E5 and 3E6 eq.) and **14** (3E6 eq.) and incubated for 4 h at RT. The solutions containing the vectors were then dialyzed against dPBS + 0.001% Pluronic to remove free molecules that had not bound to the AAV capsid.

### Titration of rAAV2 vector genomes

A total of 3 µL of AAV was treated with 20 units of DNase I (Roche #04716728001) at 37°C for 45 min to remove residual DNA in vector samples. After treatment with DNase I, 20 µL of proteinase K (20 mg/mL; MACHEREY-NAGEL # 740506) was added and the mixture incubated at 70°C for 20 min. An extraction column (NucleoSpin®RNA Virus) was then used to extract DNA from purified AAV vectors. Quantitative real time PCR (qPCR) was performed with a StepOnePlus™ Real-Time PCR System Upgrade (Life Technologies). All qPCRs were performed with a final volume of 20 µL, including primers and probes targeting the ITR2 sequence, PCR Master Mix (TaKaRa), and 5 µL of template DNA (plasmid standard or sample DNA). qPCR was carried out with an initial denaturation step at 95°C for 20 seconds, followed by 45 cycles of denaturation at 95°C for 1 second and annealing/extension at 56°C for 20 seconds. Plasmid standards were generated with seven serial dilutions (containing 108 to 102 plasmid copies), as described by D’Costa et al. [18].

### Western blot, silver staining and dot blot

All vectors were denatured at 100°C for 5 min using Laemmli sample buffer and separated by SDS-PAGE on 10% Tris-glycine polyacrylamide gels (Life Technologies). Precision Plus Protein All Blue Standards (BioRad) were used as a molecular-weight size marker. After electrophoresis, gels were either silver stained (PlusOne Silver Staining Kit, Protein; GE Healthcare) or transferred onto nitrocellulose membranes for Western blot. After transferring the proteins to nitrocellulose membrane using a transfer buffer (25 mM Tris/192 mM glycine/0.1 (w/v) SDS/20% MeOH) for 1 h at 150 mA in a Trans-Blot SD Semi-Dry Transfer Cell (Bio-Rad), the membrane was saturated for 2 h at RT with 5% semi-skimmed milk in PBS-Tween (0.1%) or with 1% gelatin, 0.1% Igepal in PBS-Tween (0.01%). After saturation, the membrane was probed with the corresponding antibody (anti-capsid polyclonal or anti-fluorescein-AP) and FITC-WF lectin (GalNAc detection) or overnight at 4°C. Three washes (15 min at RT) with PBS-Tween (0.1%) were performed between each stage to remove unbound reagents. Bands were visualized by chemiluminescence using alkaline phosphatase (AP) or horseradish peroxidase (HRP)-conjugated secondary antibodies and captured on X-ray film.

For immuno dot blot rAAV2 vectors were loaded at a dose of 10^10^ vg on a nitrocellulose paper soaked briefly in PBS prior to assembling the dot blot manifold (Bio-Rad). Nitrocellulose membrane containing the vectors was treated as for Western blotting.

The evaluation of the interactions of AAV2 particles was carried out according to GLYcoDiag’s protocol already described.[19–22] Briefly, the AAV2 particles are deposited at three concentrations (5.0 x 10^10^ particles/mL, 2.5 x 10^10^ particles/mL, 1.25 x 10^10^ particles/mL) in LEctPROFILE® plates functionalized with the following lectins: BPA *(Bauhinia purpurea)*, DBA *(Dolichos biflorus)*, WFA *(Wisteria floribunda)*, SBA *(Glycine max)*, HPA *(Helix pomatia)*, AIA *(Artocarpus intergrifolia)*, UEA-I *(Ulex europaeu*s), DSA *(Datura stramonium)* and ASGPR *(recombinant human asialoglycoprotein receptor 1)* in triplicate and incubated for 2 h at room temperature. After washing with PBS, the anti-AAV2 antibody labeled with biotin and preliminary diluted 20 times (final concentration of 2.5 ug/mL), was added to each well of the LEctPROFILE plate and incubated 30 min. Then, the plates are washed again and the extravidin-peroxidase conjugate is added for 30 min. Then, the LEctPROFILE® plates are washed with PBS and 100 μL of OPD reagent are added to each lectin wells for 15 minutes. Finally, after this last incubation, 100 uL of a 1 M HCl solution are added to stop the reaction before reading the plates with an absorbance reader at 450 nm (Fluostar OPTIMA, BMG LABTECH, France). The intensity of the signal was directly correlated with the ability of the compound to be recognized by the lectin.

### Mass spectrometry analyse of chemically modified rAAV2

Modified and unmodified rAAV2 samples were conditioned in dPBS buffer (1E12 vg/mL). Desalting and separation of protein contents were achieved on an H-Class UPLC system (Waters Corporation, Milford, USA) by injection of 10 μL of solutions onto an Acquity® CSH C18 column (2.1 mm × 100 mm, 1.7 μm, 130 Å; Waters Corporation) held at 60 °C. The mobile phase was composed of 5 % acetonitrile as solvent A and 100 % acetonitrile as solvent B, each containing 0.1 % formic acid. The elution was carried out using a gradient of solvent B in solvent A over 16 min at a constant flow rate of 300 μL/min. Mobile phase B was kept constant at 1% for 1 min, then linearly increased from 1% to 80% for 10 min, kept constant for 2 min, returned to the initial condition over 1 min, and kept constant for 2 min before the next injection. High-resolution mass spectrometry (HRMS) detection of proteins was performed by a Synapt G2 HRMS Q-TOF mass spectrometer equipped with a Z-Spray interface for electrospray ionization (Waters Corporation). The resolution mode was applied in a mass-to-charge (m/z) ratio ranging from 200 to 4,000 at a mass resolution of 25,000 Full Width Half Maximum in the positive ionization mode. Ionization parameters were as follow: capillary voltage of 3 kV, cone voltage of 30 V, desolvatation gas flow of 600 L/hr, source temperature of 120 °C, desolvatation temperature of 350 °C, Nitrogen as desolvatation gas. Data were collected in the continuum mode at a rate of four spectra per second. Leucine enkephalin solution prepared at 2 μg/mL in an acetonitrile/water (50/50, v/v) mixture was infused at a constant flow in the lock spray channel. A spectrum of 1 s was acquired every 20 s and allowed mass correction during experiments. The experimental molecular weights of proteins were finally deducted by deconvolution with the MaxEnt1 extension of MassLynx® software (version 4.1, Waters Corporation).

### Transduction of primary mouse hepatocytes

Primary mouse hepatocytes and culture medium were purchased from BIOPREDIC international (Rennes, FRANCE). Mouse hepatocytes were seeded in a 24-well plate at a density of approximately 2.5E5 cells/well. After reception, cell culture medium was removed and replaced with 1mL of basal medium (MIL600) with additives (ADD222) and incubated 2h at 37°C and 5% CO2. Primary mouse hepatocytes were transduced at MOI of 10^5^ with rAAV2 or bioconjugated rAAV2 vectors in 0.5 mL of culture medium. 6h after the transduction, 0.5 mL of fresh culture medium was added to each well. All rAAV vectors encoded for GFP. The percentage of GFP positive cells was measured by FACS analysis 72h after the transduction. Cells were dissociated with Trypsin-EDTA (Sigma-Aldrich), fixed with 4% paraformaldehyde and analysed on a BD-LSRII Flow Cytometer (BD Bioscience). All data were processed by FlowJo (V10; Flowjo LLC, Ashland, OR).

### Animal care and welfare

Experiments were performed on C57BL/6 (B6) mice. Animals were euthanized 1 month after AAV injection. Research was conducted at the UTE IRS2 (University of Nantes, France). The Institutional Animal Care and Use Committee of the Région des Pays de la Loire as well as the French Ministry for National Education, Higher Education and Research approved the protocols (authorizations #29288). Animals were randomly assigned to the different experimental groups. They were sacrificed by intravenous injection of pentobarbital sodium (Dolethal; Vetoquinol) in accordance with approved protocols.

### Intravenous injection

Mice were injected intravenously (IV) with a dose of 5E12 vg/kg of either bioconjugated rAAV2 eGFP-expressing or unmodified rAAV2 eGFP-expressing. Eight weeks old C57BL/6 (B6) mice (Charles River Laboratories) were used in this study. 9 groups were formed, 6 mice per group. Chemically modified AAV2 vectors (or formulation buffer) were administered by IV injection through the caudal tail vein, under general anaesthesia, at a dose of 5E12 vg/kg. The animals were anaesthetised with an O2/isoflurane 5% gas mixture in an induction cage, then transferred to a heating board fitted with an anaesthetic mask. The isoflurane was then set to 2%. All animals were euthanized one-month after injection, samples were collected in particular liver, spleen, muscle, heart, kidney and lung.

### Relative quantification of GFP messenger by RT-qPCR in mice tissue samples

Total RNA was extracted from snap-frozen liver, heart, skeletal muscle, lung, spleen and kidney, using Qiazol reagent (Qiagen) and treated with ezDNAse kit (Thermo Fisher Scientific), according to the manufacturer’s instructions. Then, 500 ng of this RNA was reverse transcribed using Superscript IV Vilo kit (Thermo Fisher Scientific), according to the manufacturer’s instructions. qPCR analyses were conducted on a C1000 touch thermal cycler (Bio-Rad) on 5µL of cDNA (diluted 1/20) in duplicate and premix Ex taq (Ozyme) using GFP primers and probe (Forward: 5’-AGTCCGCCCTGAGCAAAGA-3’; Reverse: 5’-GCGGTCACGAACTCCAGC-3’; Probe: 5’-CAACGAGAAGCGCGATCACATGGTC-3’). The murine Hprt1 messenger was also amplified as an endogenous control, using a primers/probe combination (Forward: 5’-TCTGTAAGAAGGATTTAAAGAGAAGCTA-3’; Reverse: 5’-ATCACATGTTTATTCCACTGAGCAA-3’; Probe, 5’-AGCTCTCGATTTCCTATCAGTAACAGC -3’). For each RNA sample, the absence of DNA contamination was confirmed by analysis of “cDNA-like samples” obtained without adding reverse transcriptase to the reaction mix. The absence of qPCR inhibition in the presence of cDNA was determined by analyzing cDNA obtained from tissue sample from a control animal, spiked with different dilutions of a standard plasmid. Results were expressed in relative quantification (RQ): RQ = 2-ΔCt = 2-(Ct GFP - Ct mHPRT1). The lower limit of quantification (LLOQ) of our test was 0.0004. GraphPad Prism 9 software was used for statistical analysis. Non-parametric Mann Whitney test was used. Samples were considered significantly different if *p ˂ 0.05, **p ˂ 0.01, ***p ˂ 0.001.

### Western blot and GFP expression

Total proteins were extracted from snap-frozen liver using RIPA buffer (Tris 10 mM pH 7.5; NaCl 150 mM; EDTA 1 mM; NP40 1%; sodium deoxycholate 0.5%; SDS 0.1%) containing protease inhibitor cocktail (Sigma-Aldrich). 25 µg of total protein extract were prepared in Laemmli buffer + 200mM final of DTT, reduced 10 min at 70°C and then loaded into a Novex 10% Tris-glycine gel (Thermo Fisher Scientific). Proteins were transferred on silica membrane using the trans-blot turbo kit (Biorad), according to the manufacturer’s instructions. Membranes were blocked in 5% skim milk, 1% NP40 (Sigma-Aldrich) in TBST (Tris-buffered saline, 0.1% Tween-20) and hybridized with an anti-GFP antibody (1:8,000, JL-8, Takarabio) and with a secondary anti-mouse IgG HRP-conjugated antibody (1:5,000, Dako P0447). For protein loading control, the same membrane was also hybridized with an anti-mouse GAPDH antibody (1:2,000, Novus IMG3073) and with a secondary anti-goat IgG HRP-conjugated antibody (1:2,000, Dako P0449). Immunoblots were visualized by ECL Chemiluminescent analysis system. Semi-quantification of GFP protein expression relative to mouse GAPDH was obtained by densitometry analysis, using the ImageQuant software.

### Immunohistochemistry GFP staining in mice

Formalin-fixed paraffin embedded liver samples were cut into 4 µm thick slices. Sections were first deparaffinized in methylcyclohexan and then rehydrated using decreasing concentrations of ethanol. Antigen unmasking was further performed through incubation in citrate buffer pH=6 (40 min, 98°C). After a permeabilization step (10 min, RT) with Triton (0.2%; VWR International, Leuven, Belgium), and a saturation step (45 min, RT) with goat serum (2%; Abcam Inc., Cambridge, MA, USA) and bovine serum albumin (5%, Sigma, St. Louis, USA), sections were then incubated (over-night, 4°C) with a mouse monoclonal antibody against GFP (1:100; JL8 clone, Thermo Fisher Scientific, Geel, Belgium). After PBS-washing, sections were incubated with an Alexa fluor 555-conjugated goat anti-murine antibody (1:300, 1 hour, RT, InVitroGen, Carlsbad, CA, USA) before washing, nuclei counterstaining with DAPI (1:1000, InVitroGen) and mounting. Whole histological preparations were digitized using fluorescence detection (Zeiss Axioscan Z1, x10 magnification). Numbers of total nuclei and positive cells based on red fluorescence were counted using QuPath software (Bankhead, P. et al. Scientific Reports, 2017 [23]) on 3 randomly selected field (2 mm2 each) enabling to observe a mean of 17882 nuclei per sample.

## Results and discussion

The successful validation of bioconjugating amino groups on AAV capsids by our research team has yielded promising *in vivo* outcomes for liver and retina targeting.[24–27] However, existing literature underscores the importance of ligand structure in its interaction with the ASGPR, knowing that the natural ligand for this receptor is galactose or *N*-acetylgalactosamine (GalNAc).[28] In a previous study, we synthesized GalNAc structure **1** (**figure 1**) and demonstrated that the density of ligands on the surface of rAAV vectors had an impact on transduction efficiency and the ability to escape neutralizing antibodies.[24] In a previous study, Mascitti *et al.* have developed a condensed and stable bicyclic bridged ketal based on GalNAc as a ligand (see sugar part of structure **2**), specifically targeting the ASGPR which exhibited almost 6-fold better affinity than GalNAc (1/Kads = 7.2 vs 40 μM). These findings underscore the significance of ligand design in enhancing the efficacy of rAAV capsid bioconjugation for targeted delivery. An X-ray crystal structures of the ASGPR bound to the bicyclic ligand, obtained by the same authors, showed that the hydroxyl group at the C6-position of the carbohydrate remains outside of the interaction site, suggesting that it could be functionalized without altering the affinity of the ligand.[29] Based on these results, we have developed novel ligands featuring a GalNAc structure as a recognition motif, a phenylisothiocyanate group at the C6 position for bioconjugation with the amino group from rAAV, as well as a triethylene glycol spacer between them to enhance overall hydrophilicity and mobility (**figure 1**).[30] Thus, compound **2** contains the bridged bicyclic ketal and the phenylisothiocyanate group at the C6 position, while compounds **3** and **4** feature a GalNAc with the phenylisothiocyanate at the C6 position and a methoxy group at either the anomeric position β or α. We opted for a methoxy group to minimize potential interactions and/or steric hindrance with the lectin pocket of ASGPR. Additionally, this choice allows us to examine the influence of the stereochemistry at the anomeric position on receptor affinity.

**Figure 1:**
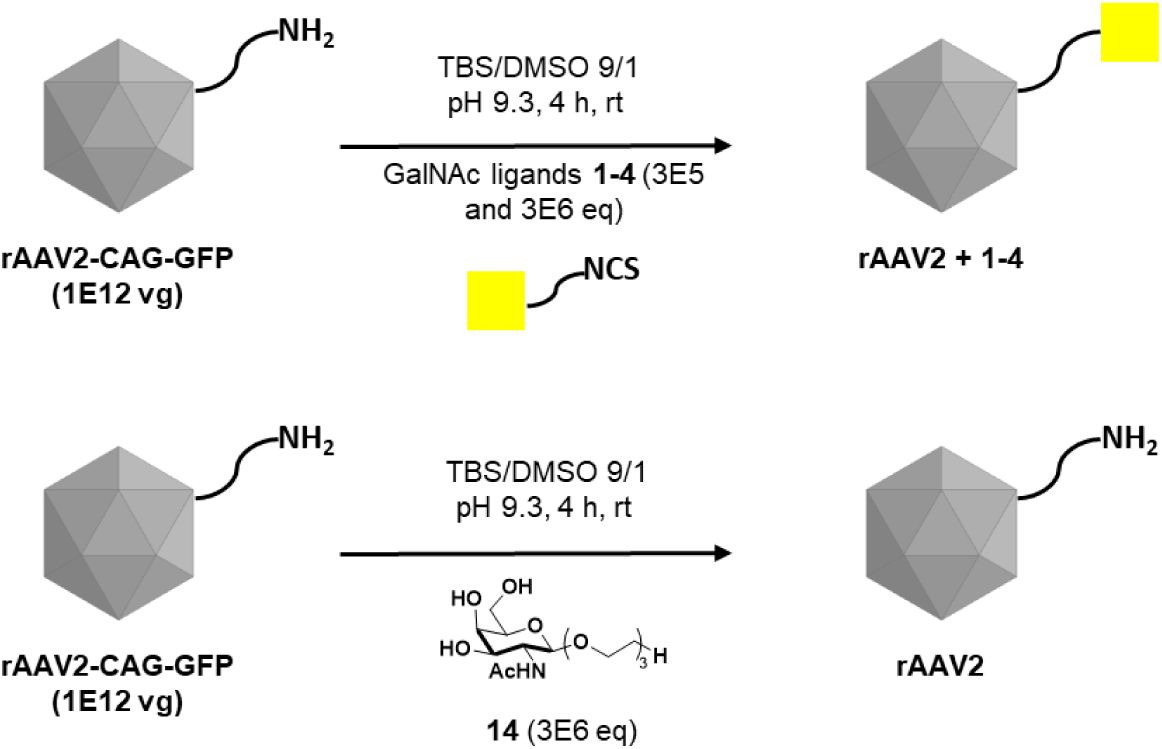
Bioconjugation step with GalNAc derivatives 1-4, 14 and rAAV2-CAG-GFP.

To access **2**, it was first possible to synthesize the key bicyclic bridged ketal intermediate **5** in 12 steps starting from the commercially available tri-*O*-acetyl-D-galactal (**Scheme 1**).[23] The hydroxyl groups at the C3- and C4-positions of the bridged GalNAc **5** were then protected in the presence of 2,2-dimethoxypropane and camphorsulfonic acid in DMF for 24 hours at 70°C to favour the formation of the thermodynamic product **6**, obtained with a yield of 93% after purification. Our initial efforts to further functionalize the hydroxyl group at the C6-position were unsuccessful using a range of organic bases in conventional *O*-alkylation conditions (not shown). Ultimately, an original procedure was performed in a biphasic mixture of DCM (for substrate solubility) and of a concentrated sodium hydroxide aqueous solution (in presence of the crown ether 15-c-5 for phase transfer), which allowed efficient access to **7**, isolated with an 80% yield. The hydroxyl groups at positions C3 and C4 were then deprotected in acidic medium, followed by palladium-catalyzed hydrogenation of the azido group in presence of PTSA which conveniently and efficiently afforded the ammonium tosylate **8**. Usage of the basic Amberlist IRN 78 resin released the corresponding primary amine, which was then reacted with bitoscanate to obtain the final bifunctional derivative **2** (**Scheme 1**).

**Figure 1:**
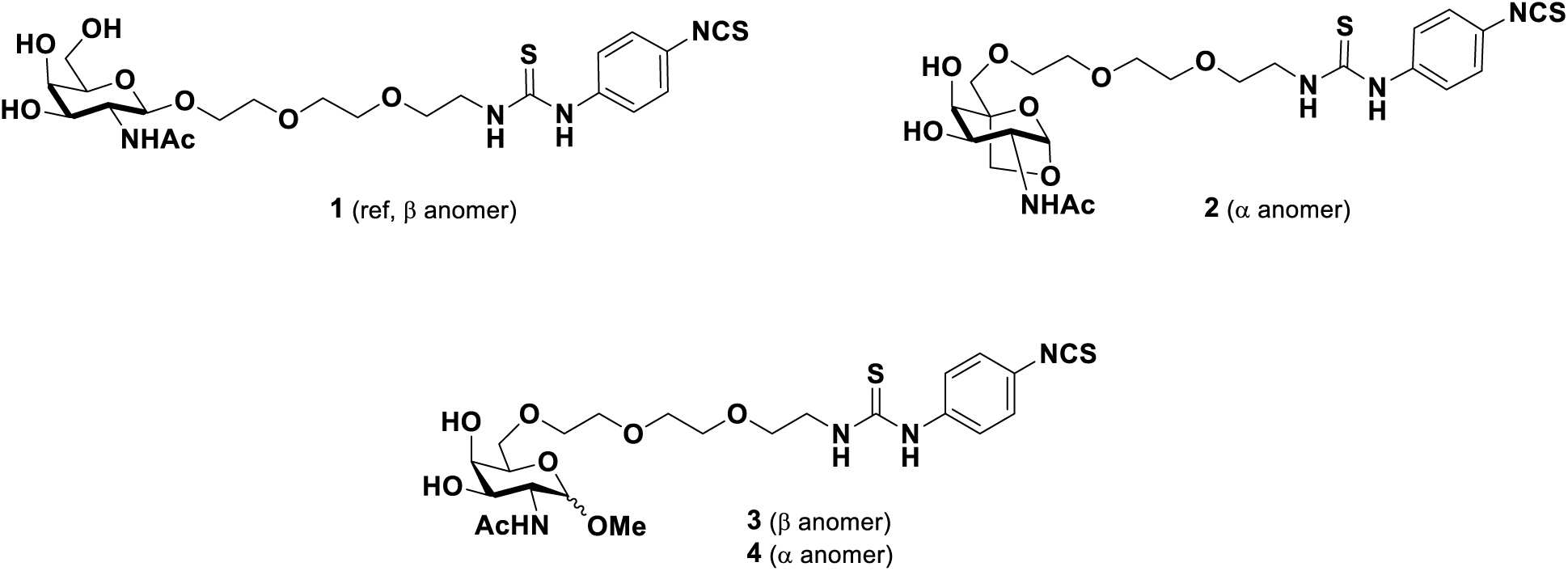
Structure of the GalNAc derivatives used for bioconjugation on rAAV2 capsid.

**Scheme 1:**
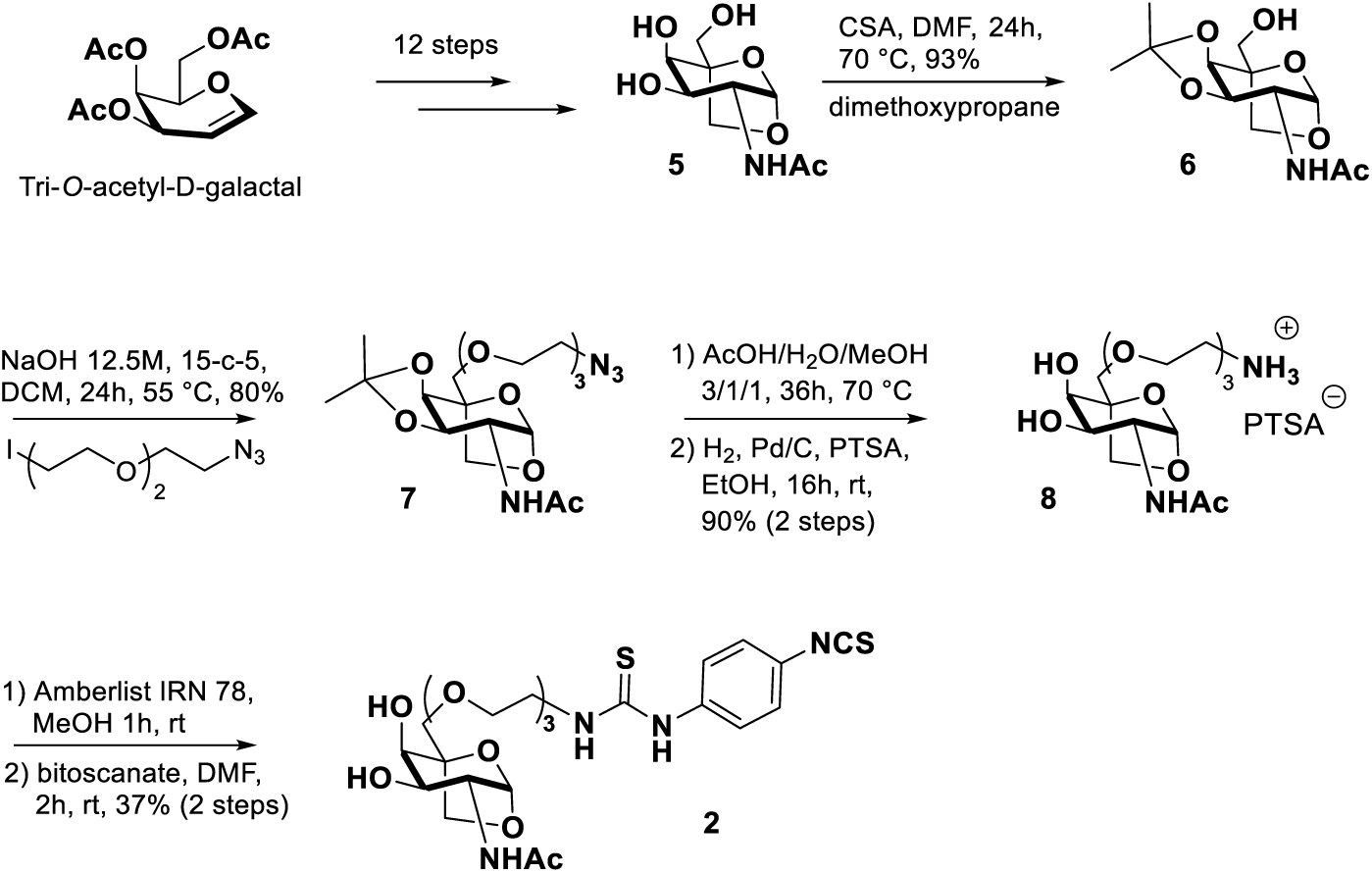
Synthesis of compound 2.

Molecules **3** and **4**, bearing the triethylene glycol chain with the aryl isothiocyanate coupling function at position C6 of GalNAc, only differ in the configuration of the methoxy group at the anomeric position. Compound **3** was synthesized in 9 steps starting from D(+)-galactosamine hydrochloride. The hydroxyl groups were firstacetylated, followed by anomeric activation to afford the corresponding glycosyl donor dihydrooxazole **9**. The latter was reacted in presence of CuCl2 in methanol to stereoselectively form the β-anomer of sugar **10**, which was isolated with a yield of 52%. After complete deactylation in basic conditions, the hydroxyl groups at positions C3 and C4 were reprotected as a cyclic ketal to give compound **11**. For the following *O*-alkylation reaction, we had to modify the reaction conditions used for the synthesis of **7**. Indeed, under the same conditions, the reaction was not complete, and an unidentified disaccharide was formed. It was found necessary to use tetrabutylammonium iodide as a phase transfer reagent and to let the reaction reflux in DCM for 72 hours. Microwave activation, changing the base (*t*BuOK, NaH, Et3N), or solvents (DMF, ACN) led to degradation of the substrate or very low conversion. Thus, compound **12** was obtained with a yield of 50% after purification by silica gel chromatography. After acetal cleavage and reduction of the azido group in previously used conditions, compound **13** (diluted in a dioxane/H2O mixture) was added in a dropwise manner to a solution of bitoscanate (excess) in dioxane, in order to prevent addition of the primary amine on both sides of bitoscanate. The final product **3** was isolated by reversed-phase C18 chromatography with a yield of 55% (**Scheme 2**).

**Scheme 2:**
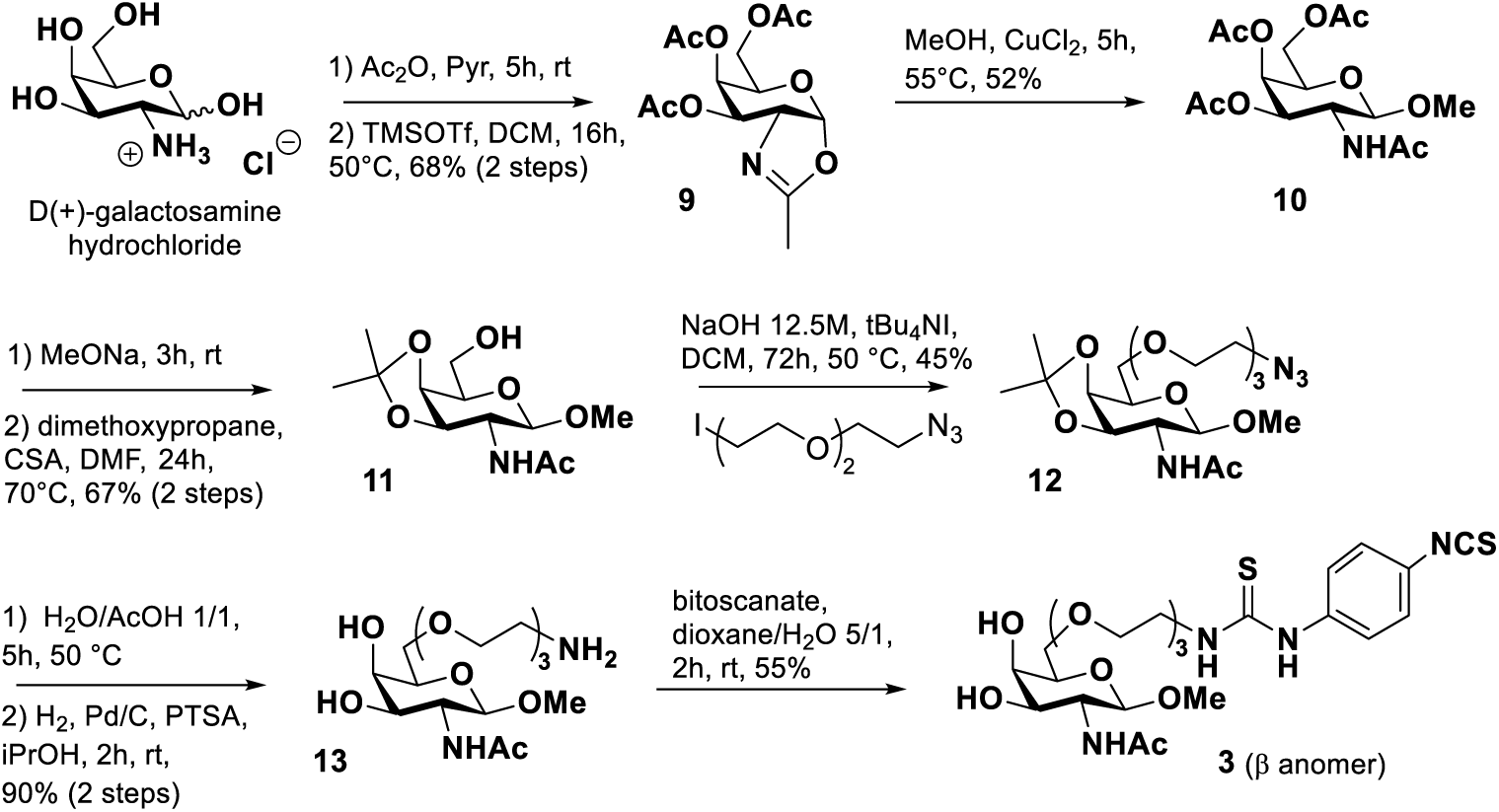
Synthesis of compound 3 (β anomer)

The compound **4** was prepared from commercially available methyl-α-D-2-acetamidogalactopyranoside, already having the desired anomeric configuration (α). In a similar way as for the β-anomer (from **10**), the envisioned sequence involves selective protection, *O-*alkylation, deprotection/reduction, and subsequent functionalization of the primary amine with bitoscanate (**Scheme S-1**). Compound **4** could be efficiently obtained in 5 steps with excellent purity.

After synthesizing the four osidic ligands, we conducted the bioconjugation step on rAAV2-CAG-GFP vectors using previously established conditions (TBS/DMSO 9/1, 4h, rt, pH 9.3).[24] The chemical coupling entailed the nucleophilic addition of the amino groups from capsid proteins to the reactive isothiocyanate function of the ligands, resulting in a stable thiourea linkage between rAAV2 and the GalNAc-derived ligands **1**-**4** (**Figure 1**).

To evaluate the impact on transduction efficiency, two different molar ratios of ligands (3E5 and 3E6) were employed to modulate the level of capsid modification using 1E12 vg total of AAV2-CAG-GFP vectors. To verify that we indeed have a covalent coupling and not ligand adsorption on the surface of the AAV capsid, we conducted the same experiment with compound **14** (3E6 eq.), which lacks the coupling function.[24] In this study, we have thus formed eight new AAVs with 4 different sugars at two concentrations.

The sugar coupling reaction on the rAAV2 surface, resulting in particles sized around 25-30 nm, underwent initial scrutiny *via* dot blot analysis (**Figure 2**). This analysis utilized the A20 antibody, which recognizes the assembled AAV2 capsid, alongside various lectins known for their selective binding to GalNAc residues (soybean (SBA), wisteria floribunda (WF), and ASGPR subunit 1).[31] The presence of positive signals, identified by the A20 antibody, confirmed the overall integrity of rAAV2 post-coupling with all ligands after subsequent dialysis. Regardless of the lectin used, no detection was observed with rAAV2 and compound **14**, lacking the NCS group, indicating the absence of physical adsorption. **Figure 2** depicted how recognition varied based on the lectins employed and the nature of the sugar chemically coupled to the rAAV2 capsid. Soybean lectin exhibited recognition of **rAAV2 + 1** under both bioconjugation conditions, albeit with low affinity for the three other sugars and only with 3E6 equivalents of ligands. In contrast, WF lectin showed strong interactions with **rAAV2 + 1**, **rAAV2 + 3**, and **rAAV2 + 4**, but weak affinity with **rAAV2 + 2**. This trend contrasted with ASGPR, where **rAAV2 + 2**, which was only slightly recognized by other lectins, was now clearly identified, showing the highest intensity with 3E6 ligand equivalents. In contrast, the **rAAV2 + 1** construct was scarcely recognized at the two concentrations used for the bioconjugation. The two other vectors (**rAAV2 + 3** and **rAAV2 + 4**) exhibited an intermediate recognition profile with this lectin and only with the 3E6 equivalents used for the coupling step. To confirm the interaction between chemically modified rAAV2 and lectins, these eight new vectors, along with an unmodified rAAV2 control and a mannose functionalized AAV2-Man(K),[26,32] as controls, were assessed against a panel of nine lectins specific to GalNAc (*Bauhinia purpurea* (BPA), *Dolichos biflorus* (DBA), *Wisteria floribunda* (WFA), *Glycine max* (SBA), *Helix pomotia* (HPA), to galactose (Gal) (*Artocarpus intergrifolia* (AIA), to fucose (Fuc) (*Ulex europeus* (UEA-I)), to GlcNAc (*Datura stramonium* (DSA), or to ASGPR, using an enzyme-linked lectin assay (ELLA) in direct binding mode with absorbance detection (**Figure S-7**).[26] Indeed, the intensity of the signal detected with each vector is directly correlated with the ability of the vector to be recognized by the lectin. The results showed no or poor recognition of (i) rAAV2 and rAAV2-Man(K) by all lectins, and (ii) **rAAV2 + 1**, **rAAV2 + 2**, **rAAV2 + 3**, **rAAV2 + 4** by Gal, GlcNAc, and Fuc lectins. Conversely, all other vectors, except those modified at both concentrations with compound **1**, exhibited interaction with ASGPr. **rAAV2 + 2** was exclusively recognized by ASGPR with the 3E5 equivalents conditions and with BPA and ASGPR for the 3E6 one, confirming its highly selective structure for the latter. It is noteworthy that **rAAV2 + 3** and **rAAV2 + 4** vectors showed no recognition by SBA and HPA lectins. This lack of recognition could be attributed to the position of the triethylene glycol chain at position 6 of the carbohydrate, likely impeding its access to the lectin pocket. These findings underscore the significance of the sugar structure coupled to the rAAV2 capsid in its interaction with different lectins and particularly the specific receptor ASGP.

**Figure 2:**
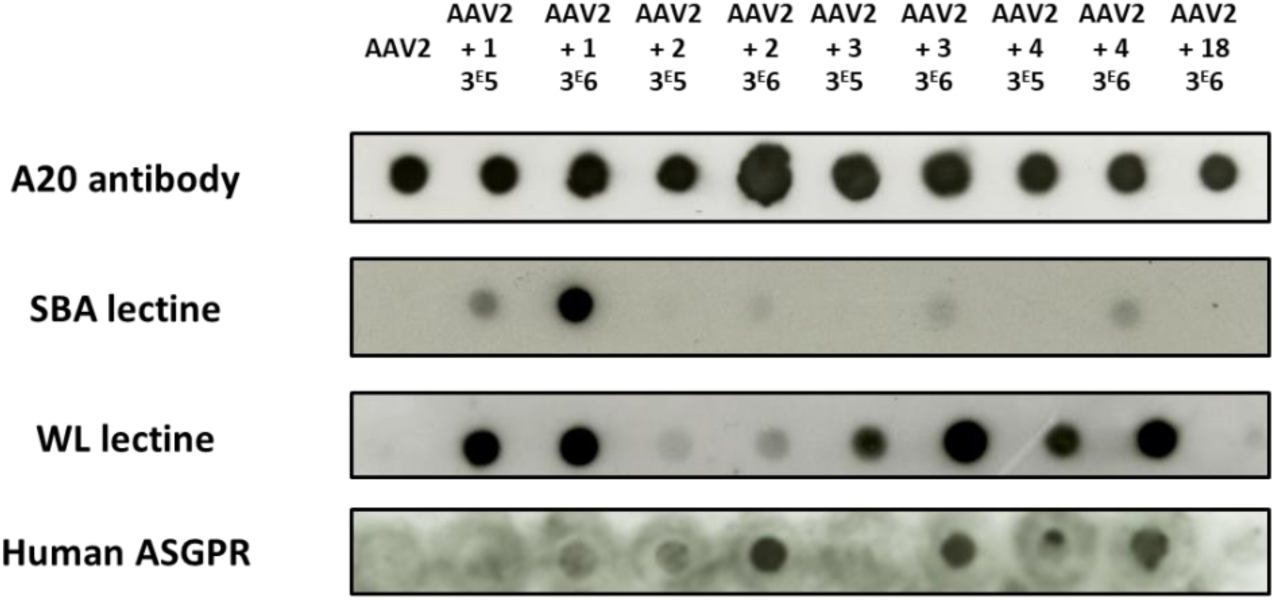
Dot blot analyses of bioconjugated rAAV vectors. 10^12^ vg of rAAV2-CAG-GFP vectors were added to a solution of **1**, **2**, **3** or **4** (3E5 and 3E6 eq.) or **18** (3E6 eq.) in TBS buffer (pH 9.3) + 10% of DMSO and incubated for 4 h at RT. 10^9^ vg of each condition was analyzed by dot blot using the A20 antibody, that recognizes the assembled capsid, SBA, WF or human ASGPR lectins.

Western blot analysis, using a capsid-specific polyclonal antibody, confirmed the integrity of VP capsid subunits regardless of molar ratios and ligands (**Figure S-8A**). Importantly, increasing the molar ratio from 3E5 to 3E6 resulted in a distinct rise in the detection of ligands on the three VPs after staining with WF lectin (**Figure S-8B**). To evaluate the purity and integrity of bioconjugated rAAV2, silver staining of various samples showed an unchanged VP1:VP2:VP3 ratio after the reaction and subsequent dialysis (**Figure S-8C**). Additionally, the molecular weight of each VP seemed to increase with escalating GalNAc derivatives loading, confirming ligand grafting onto the rAAV capsid.

To determine the number of grafts on the surface of the AAV, we used mass spectrometry to analyze the protein content of bioconjugated rAAV2. To maximize both the sensitivity and the specificity of the analysis, we primarily focused on VP3, the most abundant protein in the capsid. Results obtained for all the height chemically modified rAAV2 with 3E5 equivalents of the different ligands revealed the presence of the native VP3 protein and that in majority one ligand was bioconjugated. Upon increasing the quantity to 3E6 equivalents, we observed (*i*) the disappearance of the molecular peak of VP3, indicating complete functionalization of this protein, and (*ii*) the presence of an average of 2 ligands on each VP3 protein (**Figure S-9**). The similar behavior of the four ligands at both concentrations during bioconjugation on AAV2 demonstrates the robustness and repeatability of our technology.

All these analytical results unequivocally demonstrate that we have a covalent bioconjugation reaction on the rAAV2 with the ability to modulate the number of grafted ligands according to the experimental conditions.

Murine primary hepatocytes were transduced with all vectors at a MOI of 1E6, leveraging the targeting capability of GalNAc sugar derivatives to bind to the ASGPR, which is abundantly expressed on hepatocytes.[33] **Figure 3** illustrates that transgene (i.e. GFP) expression efficiency 48h after transduction, by FACS analyses. When compared to unmodified AAV2, no increase in GFP expression was observed for **rAAV2 + 1**, **rAAV2 + 2** and **rAAV2 + 3** modified with 3E5 eq., as well as for **rAAV2 + 3** and **rAAV2 + 4** modified with 3E6 eq. On the contrary, for **rAAV2 + 1**, **rAAV2 + 2**, and **rAAV2 + 3**, each modified with 3E6 eq., as well as **rAAV2 + 4** modified with 3E5 eq., the GFP expression was approximately two-fold higher than with unmodified rAAV2. Even if statistical significance was not reached, due to the low number of replicates, these findings underscore the maintained transduction efficiency after bioconjugation, and even more the effectiveness of some GalNAc sugar derivatives bioconjugation and modulation on rAAV2 for *in vitro* hepatocyte targeting.

**Figure 3:**
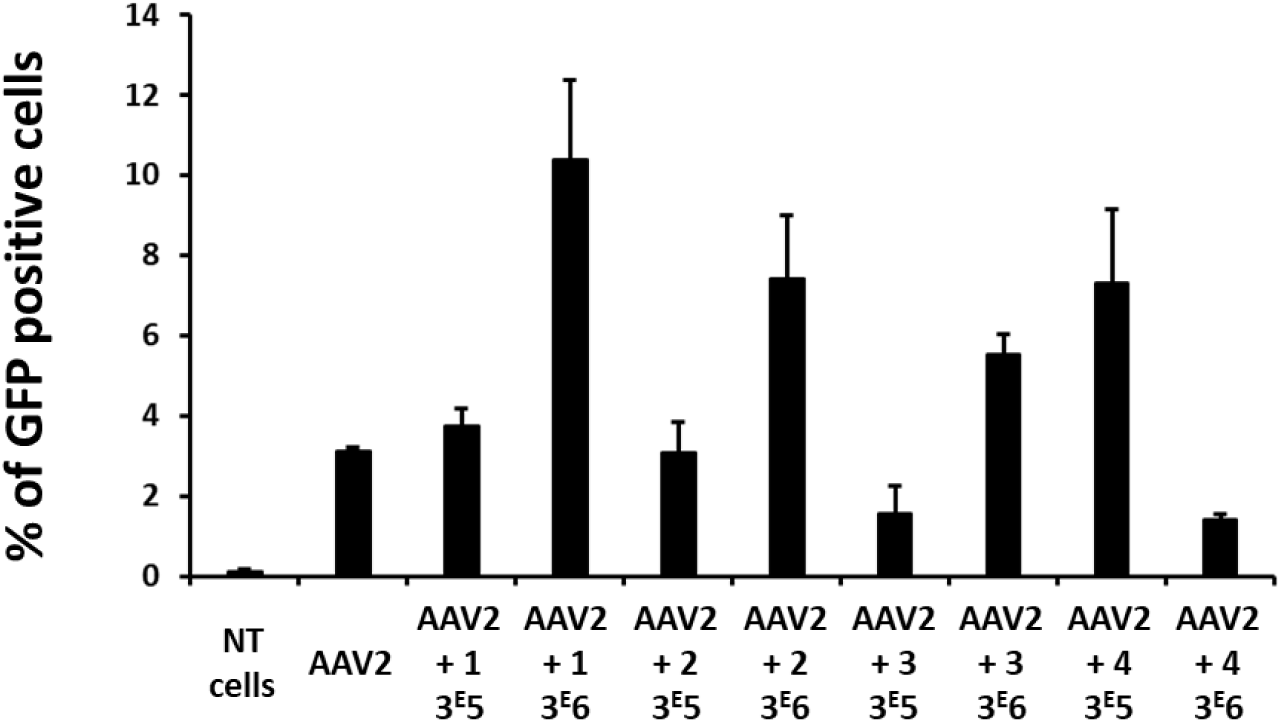
Transduction of primary mouse hepatocytes with native rAAV2 and bioconjugated rAAV2 vectors. All rAAV2 vectors encoded for GFP. The percentage of GFP positive cells was measured by FACS analysis 72 h after the transduction. Non-transduced cells (NT cells) were used as a control for the fluorescence background. Three replicates of each condition were analyzed by Kruskall Wallis test without significance. Data are represented as mean ± SD (n=3).

To evaluate the effect of the bioconjugation of GalNAc ligands **1-4** on the rAAV2 capsid lysine *in vivo*, a total of 60 healthy adult mice were intravenously injected with (*i*) the vector formulation buffer, (*ii*) the unmodified rAAV2 vector or (*iii*) the **rAAV2 + 1**, **rAAV2 + 2**, **rAAV2 + 3** and **rAAV2 + 4** vectors, each modified with 3E5 and 3E6 equivalents (n = 6 per group). All vectors were administered at a single dose of 5E12 vg/kg and carried the eGFP transgene under the control of a CAG promoter. All animals were euthanized at 1-month post-injection to collect liver, heart, spleen, skeletal muscle (quadriceps), lung and kidney for post-mortem analysis.

To compare the gene transfer efficiency in the liver between the unmodified rAAV2 vector and all the **rAAV2 + 1**, **rAAV2 + 2**, **rAAV2 + 3** and **rAAV2 + 4** vectors, the levels of eGFP RNA transcript and protein, expressed from the genome of the different injected vectors, were quantified. As expected, relative eGFP messenger quantification (Rq) by RT-qPCR (**Figure 4A**) showed no signal in mice injected with the vector formulation buffer, whereas eGFP messenger was detected in the group injected with the unmodified rAAV2, with a Rq mean of 0.2697. For mice injected with the modified vectors, a significant increase in GFP messenger level is observed for the **rAAV2 + 3** (3E6 eq.) and the **rAAV2 + 4** (3E5 eq.) groups, as observed *in vitro*, with approximately 6- and 4-fold higher Rq values respectively compared to the unmodified rAAV2 group. For animals injected with **rAAV2 + 1** (3E5 and 3E6 eq.) and **rAAV2 + 2** (3E6 eq.) vectors, GFP expression is similar to that of the unmodified AAV2 group. Finally, for the **rAAV2 + 2** (3E6 eq.), **rAAV2 + 3** (3E5 eq.) and **rAAV2 + 4** (3E6 eq.) groups, the Rq mean values were respectively 0.0008, 0.0246 and 0.0007, just very close to the LLOQ (lower limit of quantification) of 0.0004.

**Figure 4:**
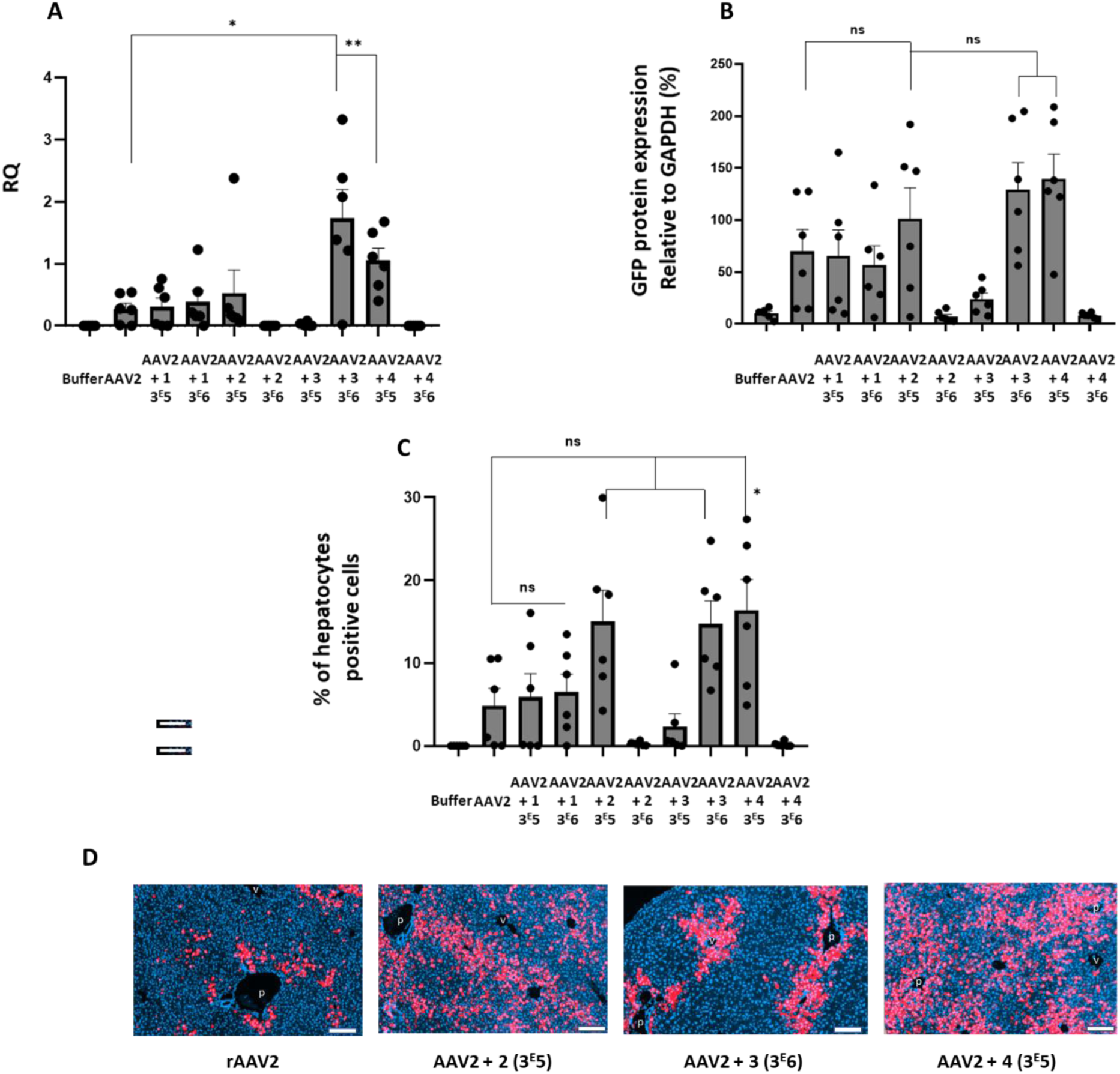
A) Relative quantification of *GFP* mRNA levels (relative to rat *Hprt1* mRNA levels) analyzed by RTqPCR in liver 4 weeks post injection in mice injected with vehicle (buffer), rAAV2, or bioconjugated rAAV2 vectors. Results are expressed as mean ± SEM. The Mann-Whitney U-test was used for statistical analyses: **p<0.01, *p<0.1. Lower limit of quantification (LLOQ = 0.0004. B) Levels of GFP protein expression in mouse liver detected by Western blot 1 month after injection. C) Percentage of GFP-positive cells in liver, detected by GFP immunolabelling. D) Representative image of liver after GFP immunolabelling (red). p, portal vein branches; c, centro-lobular veins. DAPI counter-staining of nuclei is shown in blue. Scale bars=200 µm.

The relative eGFP protein semi-quantification by western blot (**Figure 4B, S-10**) supports these results. Indeed, **rAAV2 + 2** (3E6 eq.), **rAAV2 + 3** (3E5 eq.) and **rAAV2 + 4** (3E6 eq.) showed lower eGFP expression levels than the unmodified AAV2 group, and **rAAV2 + 1** (3E5 and 3E6 eq.) groups showed similar eGFP expression levels as the unmodified AAV2 group. On the contrary, eGFP expression levels were found about 1.5-fold higher in the **rAAV2 + 2** (3E5 eq.) group and 2-fold higher in the **rAAV2 + 3** (3E6 eq.) and **rAAV2 + 4** (3E5 eq.) groups, when compared to the unmodified rAAV2 group. Immunohistochemistry analysis shows higher percentages of GFP-positive cells in the livers of mice injected with **rAAV2 + 2** (3E5 eq.), **rAAV2 + 3** (3E6 eq.) and **rAAV2 + 4** (3E5 eq.) compared to the unmodified rAAV2 (mean of 15%, 15%, 16% and 5% respectively) (**Figure 4C**). GFP-positive hepatocytes (in red) were observed in the liver parenchyma of animals injected with rAAV2, mainly distributed as clusters close to portal tracts, with sparse GFP-positive cells detected around centro-lobular veins. By contrast, mice injected with **rAAV2 + 2** (3E5 eq.), **rAAV2 + 3** (3E6 eq.) and **rAAV2 + 4** (3E5 eq.) showed a broader distribution of GFP-positive cells, which were clustered around portal tracts and also detected throughout the liver parenchyma and in greater numbers around the portal vein (**Figure 4D**). Considering their similar appearance (polygonal shape with a round nucleus) and their inclusion into Remack trabeculae of the liver parenchyma, the vast majority of GFP-positive cells were identified as hepatocytes (in red on **Figure 4D**). Following injection with rAAV2, GFP-positive hepatocytes were mainly distributed as clusters close to portal tracts, with sparse GFP-positive cells detected around centro-lobular veins. By contrast, mice injected with **rAAV2 + 2** (3E5 eq.), **rAAV2 + 3** (3E6 eq.) and **rAAV2 + 4** (3E5 eq.) showed a broader distribution of GFP-positive cells, which were not only densively clustered around portal tracts but also detected isolated throughout the liver parenchyma and in small groups around the portal vein (**Figure 4D**).

Finally, the level of eGFP RNA transcript was quantified in non-targeted tissues (*i.e*. heart, quadriceps muscle, lung, spleen, and kidney). In all organs, Rq values were low and similar between unmodified and modified rAAV2 groups. These data showed that, at the tested dose, bioconjugation of rAAV2 does not increase the transduction of these non-targeted tissues (**Table S1**).

In summary, these findings indicate that *in vivo* liver gene transfer is significantly more efficient with rAAV2 + **2** (3E5 eq.), rAAV2 + **3** (3E6 eq.), and rAAV2 + **4** (3E5 eq.) vectors compared to the unmodified rAAV2 vector, as evidenced by higher mRNA expression levels, protein expression levels, and the percentage of transduced hepatocytes, along with the absence of off-target effects. These results, achieved with an injected dose of 5E12 vg/kg of rAAV2, are particularly encouraging given that this dose is intermediate compared to the 1E13 vg/kg dose used by Greig *et al*..[34] Additionally, our results highlight the critical role of the selected sugar structure for the bioconjugation step and its modulation on the capsid surface, especially for rAAV2. Although rAAV2 is not the most effective serotype for this application, it clearly demonstrates the beneficial effects of our technology.

However, further studies are needed to identify a lead structure and optimal bioconjugation conditions to ensure maximal transduction efficiency and targeting. Future research will focus on intracellular trafficking studies, from viral vector entry to final protein expression, to better understand these results. This will facilitate the development of the most efficient chemically modified vector in terms of ligand structure, bioconjugation conditions, and serotype for potential clinical applications.

## Conclusion

Enhancing vector transduction efficiency stands as a critical imperative in surmounting the myriad challenges inherent in rAAV gene therapy. The bioconjugation approach directly on the capsid offers promising avenues for bolstering targeting precision, optimizing cellular entry, and increasing expression levels of the intended protein. Through an integrated approach melding chemistry with vectorology, our investigation underscores the pivotal role played by both the structure and quantity of grafted molecules onto the lysine residues of the native rAAV2 capsid. Our findings unveil discernible impacts on vector transduction efficacy, gene expression, and hepatocyte population percentages, both *in vitro* and *in vivo* within the liver, the focal point of our study. Chemical refinement of therapeutic viruses presents a compelling alternative to genetic manipulation techniques for enhancing organ tropism and protein expression within target cells. Future endeavors will entail the evaluation of this technology with therapeutic transgenes tailored for liver-specific applications and its extension across diverse serotypes. Additionally, we intend to embark on intracellular investigations to elucidate the trafficking mechanisms governing bioconjugated rAAV vectors, paving the way for the conception of even more efficacious treatments.

## Acknowledgments

The authors thank ViVeM of TaRGeT, UMR 1089 (CPV, INSERM and Nantes Université, which is a bioproduction and biotherapy national integrator (ANR-22-AIBB-0001), http://umr1089.univ-nantes.fr) for the production of the rAAV vectors used in this study.

This research was supported by the Fondation d’Entreprise Thérapie Génique en Pays de Loire, the Centre Hospitalier Universitaire (CHU) of Nantes, the Institut National de la Santé et de la Recherche Médicale (INSERM), Nantes Université, and by grants from the French National Agency for Research (“Investissements d’Avenir” Equipex ArronaxPlus.n°ANR-11-EQPX-0004 and ChemAAV (ANR-19-CE18-0001)).

## CRediT authorship contribution statement

**Pierre-Alban Lalys** (Conceptualization; Methodology), **Audrey Bourdon** (Conceptualization; Methodology; Supervision; Validation; Writing – original draft; Writing – review & editing), **Mohammed Bouzelha** (Conceptualization; Methodology; Validation), **Dimitri Alvarez-Dorta** (Methodology; Validation), **Karine Pavageau** (Methodology; Validation), **Tiphaine Girard** (Methodology), **Roxanne Peumery** (Methodology), **Zakaria Bouchouireb** (Methodology), **Maia Marchand** (Methodology), **Anthony Mellet** (Methodology), **Mireille Ledevin** (Formal analysis; Methodology), **Sébastien Depienne** (Visualization; Writing – review & editing), **Mickaël Guilbaud** (Conceptualization; Formal analysis; Methodology; Writing – review & editing), **Mikaël Croyal** (Conceptualization; Formal analysis; Methodology; Writing – review & editing), **Sébastien G. Gouin** (Conceptualization; Methodology; Writing – review & editing), **Benoît Roubinet** (Lectins interaction analysis), **Ludovic Landemarre** (Lectins interaction analysis, Resources), **Oumeya Adjali** (Funding acquisition; Project administration; Resources), **Eduard Ayuso** (Conceptualization; Funding acquisition; Supervision), **Thibaut Larcher** (Conceptualization; Methodology; Supervision; Writing – original draft), **Caroline Le Guiner** (Supervision; Validation; Writing – original draft; Writing – review & editing), **Mathieu Mével** (Conceptualization; Funding acquisition; Supervision; Validation; Writing – original draft; Writing – review & editing), **David Deniaud** (Conceptualization; Supervision; Funding acquisition; Methodology; Resources; Validation; Writing – original draft; Writing – review & editing).

## Data availability

All relevant data are available from the authors on request and are included within the manuscript.

## Appendix A. Supplementary data

Synthesis, ^1^H NMR, ^13^C NMR, bioconjugaison characterization, ELLA assay, mass spectrometry analyses, transduction of primary hepatocytes, western blot of total proteins, mRNA GFP quantification.

